# Turning plants from passive to active material: FERONIA and microtubules independently contribute to mechanical feedback

**DOI:** 10.1101/2021.03.24.436809

**Authors:** Alice Malivert, Özer Erguvan, Antoine Chevallier, Antoine Dehem, Rodrigue Friaud, Mengying Liu, Marjolaine Martin, Théophile Peyraud, Olivier Hamant, Stéphane Verger

## Abstract

To survive, cells must constantly resist mechanical stress. In plants, this involves the reinforcement of cell walls, notably through microtubule-dependent cellulose deposition, and thus wall sensing. Several receptor-like kinases have been proposed to act as mechanosensors. Here we tested whether the microtubule response to stress acts downstream of known wall sensors. Using a multi-step screen with eleven mutant lines, we identify FERONIA as the primary candidate for controlling the microtubule response to stress. However, when performing mechanical perturbations, we show that the microtubule response to stress can be independent from FER. We reveal that the *feronia* phenotype can be partially rescued by reducing tensile stress levels. Conversely, in the absence of both microtubules and FER, cells swell and burst like soap bubbles. Altogether, this shows that the microtubule response to stress acts as an independent pathway to resist stress, in parallel to FER. We propose that both pathways are key components to turn plant cells from passive to active material.

## Introduction

All living organisms use mechanical forces as instructive cues during their development (Wolfenson et al., 2019)(Hamant and Saunders, 2020). They also share a common mechanical property: cells are pressurized by osmotic pressure and thus experience cortical tension. Osmotic pressure in plants is several orders of magnitude higher than that of animal cells, and it is counterbalanced by stiff cell walls (Schopfer, 2006). Regulating the mechanical properties of cell walls, through the perception of wall tension and integrity, is thus crucial for plant growth and development (Bacete and Hamann, 2020)(Trinh et al., 2021).

Plant cell walls are composed of load-bearing cellulose microfibrils, tethered by a matrix made of polysaccharides and structural proteins (Cosgrove, 2005)(Anderson and Kieber, 2020). The deposition of cellulose microfibrils is generally guided by cortical microtubules (Paredez et al., 2006). Beyond the average stiffness, the orientation of cellulose microfibrils controls the mechanical anisotropy of the wall. There is now ample evidence showing that cortical microtubules align with maximal tensile stress directions (Green and King, 1966)(Williamson, 1990)(Hejnowicz et al., 2000)(Hamant et al., 2008)(Verger et al., 2018). This provides a feedback loop in which shape and growth, whether at the individual cell or whole organ scale, prescribes a pattern of stress, to which cells resist by reinforcing their walls along maximal tensile stress directions (Hamant et al., 2008)(Sampathkumar et al., 2014).

Although they are significantly less stiff, other wall components contribute to wall properties. In particular, pectins can partially rescue defects in cellulose synthesis in young cell walls. For instance, isoxaben treatment, which inhibits cellulose deposition through the internalization of CESA complexes, leads to thicker walls that are enriched in pectin (Manfield et al., 2004). Similarly, in young hypocotyls, pectin polarities precede the formation of mechanically anisotropic walls (Peaucelle et al., 2015). In contrast to cellulose deposition, pectin, as well as all other matrix components, are secreted to the cell wall (Cosgrove, 2005)(Anderson and Kieber, 2020). Therefore, in principle this provides an alternative mechanism for the cell to resist wall tension or damage. As for microtubules, the related mechanotransduction pathway is largely unknown. Yet, over the past decade, *Catharanthus roseus* Receptor-like kinases (CrRLKs) have emerged as key players. Although the link with mechanical stress remains to be formally established, THESEUS1 (THE1) has been involved in the wall integrity pathway (Hématy et al., 2007)(Gonneau et al., 2018)(Engelsdorf et al., 2018). Based on defective root growth behaviour on stiff interface, calcium signalling, pH response, and *TOUCH* gene expression, FERONIA (FER) acts as a mechanosensor (Shih et al., 2014). FER can sense the status of the cell wall, notably when salinity rises, through pectin binding (Feng et al., 2018). Last, it was recently proposed that FER also monitors microtubule behaviour through a cascade involving Rho GTPases (ROP6) and the microtubule severing protein katanin (Lin et al., 2018). Here, through a reverse genetic screen on wall sensors and using a suite of mechanical tests, we show that our best wall sensor candidate FER is not required for the microtubule response to stress, further suggesting that the microtubule response to stress can be more autonomous than anticipated. We also reveal that FER-dependent wall integrity pathway depends on wall tension, and that both FER and the microtubule response to stress contribute to wall integrity.

## Results

### Altered pavement cell shape as a proxy for defective response to mechanical stress

We first used a morphometric proxy to test the involvement of wall sensors in the microtubule response to stress. The shape of Arabidopsis pavement cells has been proposed to be actively maintained and amplified by the microtubule response to mechanical stress. Indeed, necks in such cells prescribe highly anisotropic tensile stresses locally, to which microtubule arrays and thus cellulose deposition align (Sampathkumar et al., 2014)(Sapala et al., 2018)(Bidhendi et al., 2019). We reasoned that the shape of pavement cells could be used in a mutant screen as a proxy for defects in that mechanical feedback loop. So far, such screens have been focused on either the intracellular biochemical cues behind cell-cell coordination (Fu et al., 2005) or the contribution of cell wall properties in cell shape (Majda et al., 2017). Whether wall sensors are involved in pavement cell shape remains ill-described. Here we focused on mutants impaired in receptor-like kinases that are highly expressed in the epidermis and aerial parts of the plant during early development and that exhibit an established link with the cell wall (and their closest homologs), namely *feronia (fer), theseus1 (the1)*, *theseus1/feronia-related1* (*tfr1*; *at5g24010)*, *curvy1* (*cvy1)*, *hercules receptor kinase 1* (*herk1)*, *herk2, mdis1-interacting receptor-like kinase2 (mik2-1), wall-associated kinase 1 (wak1)*, *wak2*, *wak3*, *wak4* (see Supplementary Table 1). We imaged and quantified the pavement cell shapes in receptor-like kinase candidate mutants, with the aim to select the ones with the strongest cell shape defects (Figure 1A).

**Figure 1.**
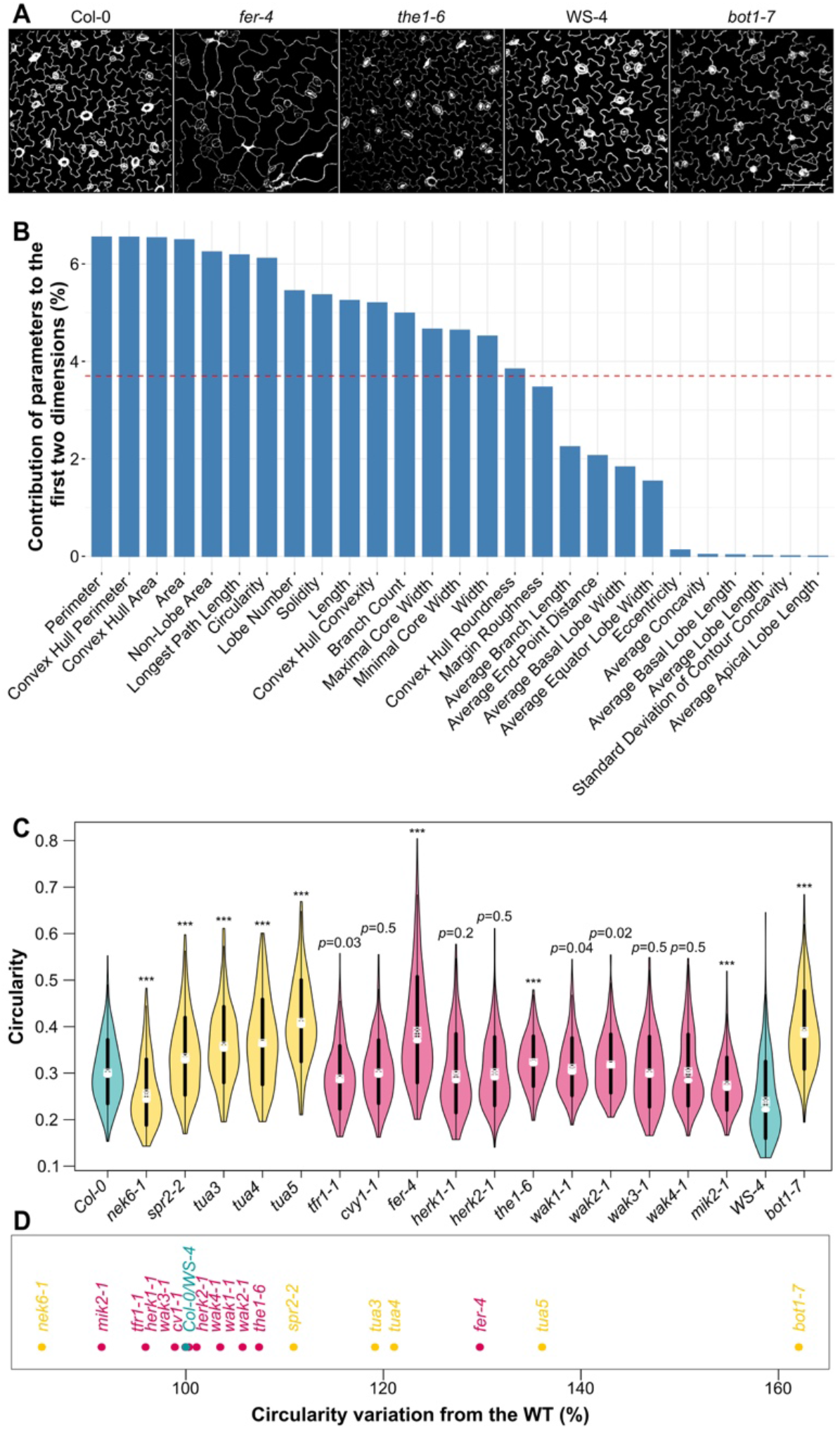
Pavement cell shape as a proxy for microtubule response to mechanical stress in receptor-like kinase mutants **A**. Representative images of *Col-0, WS-4, fer-4, the1-6* and *bot1-7* pavement cells. Samples were PI stained and cell contours were extracted with MorphoGraphX and projected in 2D. Scale=100μm. **B**. Relative contribution of 27 shape descriptors to pavement cell shape, as assessed by Principal Component Analysis. **C.** Circularity (violin plots) of pavement cells and *p*-values (*p*) of Dunn tests for the WT (in blue, *Col-0, WS-4*), for the microtubule regulator (in orange, *nek6-1, spr2-2, tua3, tua4, tua5, bot1-7*) and the receptor-like kinase mutants (in pink, *tfr1-1, cvy1-1, fer-4, herk1-1, herk2-1, the1-4, the1-6, wak1-1, wak2-1, wak3-1, wak4-1, mik2-1*). **D**. Percentage of increase or decrease in pavement cell circularity from the WT (in blue, *Col-0, WS-4*), for the microtubule regulator (in orange, *nek6-1, spr2-2, tua3, tua4, tua5, bot1-7*) and the receptor-like kinase mutants (in pink, *tfr1-1, cvy1-1, fer-4, herk1-1, herk2-1, the1-4, the1-6, wak1-1, wak2-1, wak3-1, wak4-1, mik2-1*)

To do so, we extracted the epidermal signal and used PaCeQuant to segment pavement cells and measure 27 shape descriptors (see Material and methods, (Möller et al., 2017) (Supplementary Figure 1A)). To select the most discriminating PaCeQuant shape descriptor, we first performed a principal component analysis on our data. We compared the contribution of each parameter to the two axes with the most variation (Figure 1B). The cell perimeter was first, followed by the convex hull perimeter, the convex hull area, the cell area, the non-lobe area and the longest path length, then by circularity (for relevant definition, see Supplementary Figure 1B).

To confirm that such shape descriptors are pertinent, and knowing that cortical microtubules are well-known pavement cell shape regulators, we used mutants with microtubule defects as positive controls. We performed the same analysis on mutants with a reported enhanced microtubule response to stress (*nek6* (Takatani et al., 2020)) and a reported reduced response to stress (*bot1* (Uyttewaal et al., 2012). We also included lines with tubulin mutations affecting microtubule dynamics (*tua3^D205N^, tua4^S178Δ^*, and *tua6^A281T^* referred as *tua3*, *tua4* and *tua5* in the following (Ishida et al., 2007)(Matsumoto et al., 2010)) and *spr2* with a reported enhanced cortical microtubule response to stress (Hervieux et al., 2016) but also ambivalent regulatory role in microtubule severing depending on tissue (Wightman et al., 2013)(Fan et al., 2018)(Nakamura et al., 2018). We found that cell perimeter, convex hull perimeter, convex hull area, cell area, and the longest path length descriptors were not sufficient to discriminate the pavement cell shape phenotype of those microtubules regulators (cell perimeter: *p_spr2-2_*=0.07; *p_tua3_*=0.018; *p_tua4_*=0.4; convex hull perimeter: *p_tua3_*=0.39; *p_tua5_*=0.011; convex hull area: *p_tua3_*=0.5; *p_tua5_*=0.03; cell area: *p_tua3_*=0.06; *p_tua5_*=0.07; longest path length: *p_tua3_*=0.28; *p_tua4_*=0.1 n_*spr2-2*_=262; n_*tua3*_=204; n_*tua4*_=230; n_*tua5*_=297; n_Col-0_=428). In contrast, all microtubule regulator mutant lines tested exhibited a defect in non-lobe area and circularity (non-lobe area: *p_nek6-1_*=0.009; *p_spr2-2_*<10^−3^; *p_tua3_*=0.003; *p_tua4_*<10^−3^; *p_tua5_*<10^−3^; *p_bot1-7_*<10^−3^; circularity: *p_nek6-1_*<10^−3^; *p_spr2-2_*<10^−3^; *p_tua3_*<10^−3^; *p_tua4_*<10^−3^; *p_tua5_*<10^−3^; *p_bot1-7_*<10^−3^, n_*nek6-1*_=183; n_*spr2-2*_=262; n_*tua3*_=204; n_*tua4*_=230; n_*tua5*_=297; n_*bot1-7*_=252; n_Col-0_=428; n_WS-4_=294, Figure 1C and Supplementary Figure 1C).

9 of the 11 receptor-like kinase mutants exhibited a non-lobe area significantly different than that of the WT. *tfr1-1*, *fer-4*, *herk1-1*, *herk2-1*, and *wak4-1* displayed a non-lobe area significantly higher than that of the WT (*p_tfr1-1_*<10^−3^; *p_fer-4_*<10^−3^; *p_herk1-1_*<10^−3^; *p_herk2-1_*<10^−3^; *p_wak4-1_*=0.007; n_*tfr1-1*_=244; n_*fer-4*_=321; n_*herk1-1*_=225; n_*herk2-1*_=238; n_*wak4-1*_=204; n_*Col-0*_=428, Supplementary Figure 1C) while the non-lobe area of *cvy1-1*, *the1-6*, *wak1-1* and *mik2-1* was significantly lower than that of the WT (*p_cvy1-1_*<10^−3^; *p_the1-6_*<10^−3^; *p_wak1-1_*<10^−3^; *p_mik2-1_*<10^−3^; n_*cvy1-1*_=277; n_*the1-6*_=174; n_*wak1-1*_=295; n_*mik2-1*_=177; n_*Col-0*_=428). Only *wak2-*1 and *wak3-1* displayed non-lobe area values that were non-significantly different from that of the WT (*p_wak2-1_*=0.1; *p_wak3-1_*=0.03; n_*wak2-1*_=271; n_*wak3-1*_=202; n_Col-0_=428). Thus, non-lobe area is not a discriminant parameter in our screen. We decided to study the next most variable parameter with defects in known microtubule regulator lines – circularity – for the rest of the analysis, justifying *a posteriori* a common choice in the literature on pavement cell shape (Zhang et al., 2011).

Among the receptor-like kinase mutant tested, the pavement cells in *fer-4*, *the1-6*, *wak2-1* and *mik2-1* were significantly more circular than the WT supporting the hypothesis that the corresponding proteins could contribute to the microtubule response to stress in pavement cells (*p_fer-4_*<10^−3^; *p_the1-6_*<10^−3^; *p_wak2-1_*=0.002; *p_mik2-1_*<10^−3^; n_*fer-4*_=321; n_*the1-6*_=174; n_*wak2-1*_=271; n_*mik2-1*_=177; n_Col-0_=428, Figure 1C). *wak1-1* and *tfr1-1* also exhibited increased pavement cell circularity, albeit much less significantly (*p_wak1-1_*=0.04; *p_tfr1-1_*=0.03; n_*wak1-1*_=295; n_*tfr1-1*_=244). Pavement cells in all the other receptor-like kinase mutants (*cvy1-1*, *herk1-1*; *herk2-1*; *wak3-1*; *wak4-1*) displayed a circularity comparable to that of the WT (p_*cvy1-1*_=0.49; *p_herk1-1_*=0.2; *p_herk2-1_*=0.47; *p_wak3-1_*=0.47; *p_wak4-1_*=0.5; n_*cvy1-1*_=277; n_*herk1-1*_=225; n_*herk2-1*_=238; n_*wak3-1*_=202; n_*wak4-1*_=204; n_Col-0_=428). To distinguish the relative contributions of the most affected mutants, we quantified the deviation of circularity from the WT. Among all the receptor-like kinases tested, the candidate mutants with the largest defect in circularity when compared to the WT were *fer-4* and, to a lesser extent, *the1-6* (Figure 1D). *fer-4* cells were 30 % more circular and *the1-6* cells were 7% more circular than the WT, values that were comparable to that of microtubule regulator mutants, such as *spr2-2* or *bot1-7* (Figure 1D).

Note that the same trend was obtained when considering solidity, another parameter often used to characterize lobe formation (Vőfély et al., 2019) (Supplementary Figure 1D). Altogether, these data suggest that *fer-4* may have the most significant defect in microtubule dynamics hinting to a potential role of FERONIA in the microtubule response to mechanical stress.

### Differential response of receptor-like kinase mutants to isoxaben

To challenge the results from this initial screen, we next used a well-established protocol to mechanically perturb cell walls. Isoxaben inhibits the biosynthesis of cellulose (Scheible et al., 2001), and thus weakens the wall. In past work, such treatment were shown to induce a hyper-alignment of cortical microtubules at the shoot apical meristem and in cotyledon pavement cells (Heisler et al., 2010)(Sampathkumar et al., 2014), consistent with a response to increased tensile stress levels in the cell wall. Note that isoxaben can also trigger other responses, including ROS production, lignification, and changes in gene expression (Hématy et al., 2007). Thus, depending on time and dose, isoxaben may also ultimately reduce stress level (Engelsdorf et al., 2019).

We grew the seedlings in a medium containing 1 nM isoxaben or the same volume of DMSO, in the dark (Figure 2A). We then measured the length of the etiolated hypocotyls four days after germination. After isoxaben treatment, four-day old WT seedlings exhibited a shorter hypocotyl (by 42% for Col-0, 34% for WS-4, n_Col-0 DMSO_=287, n_Col-0 iso_=271, n_WS-4 DMSO_=101, n_WS-4 iso_=96). To compare WT and mutants, we normalized the obtained distribution of lengths to the same mean and standard deviation as the control, thus providing a hypocotyl length index (Figure 2B, Supplementary Figure 2A). Treated mutants with a relatively shorter hypocotyl than the treated WT were labelled more sensitive to isoxaben whereas treated mutants with a relatively longer hypocotyl than the treated WT were labelled less sensitive than the WT.

**Figure 2.**
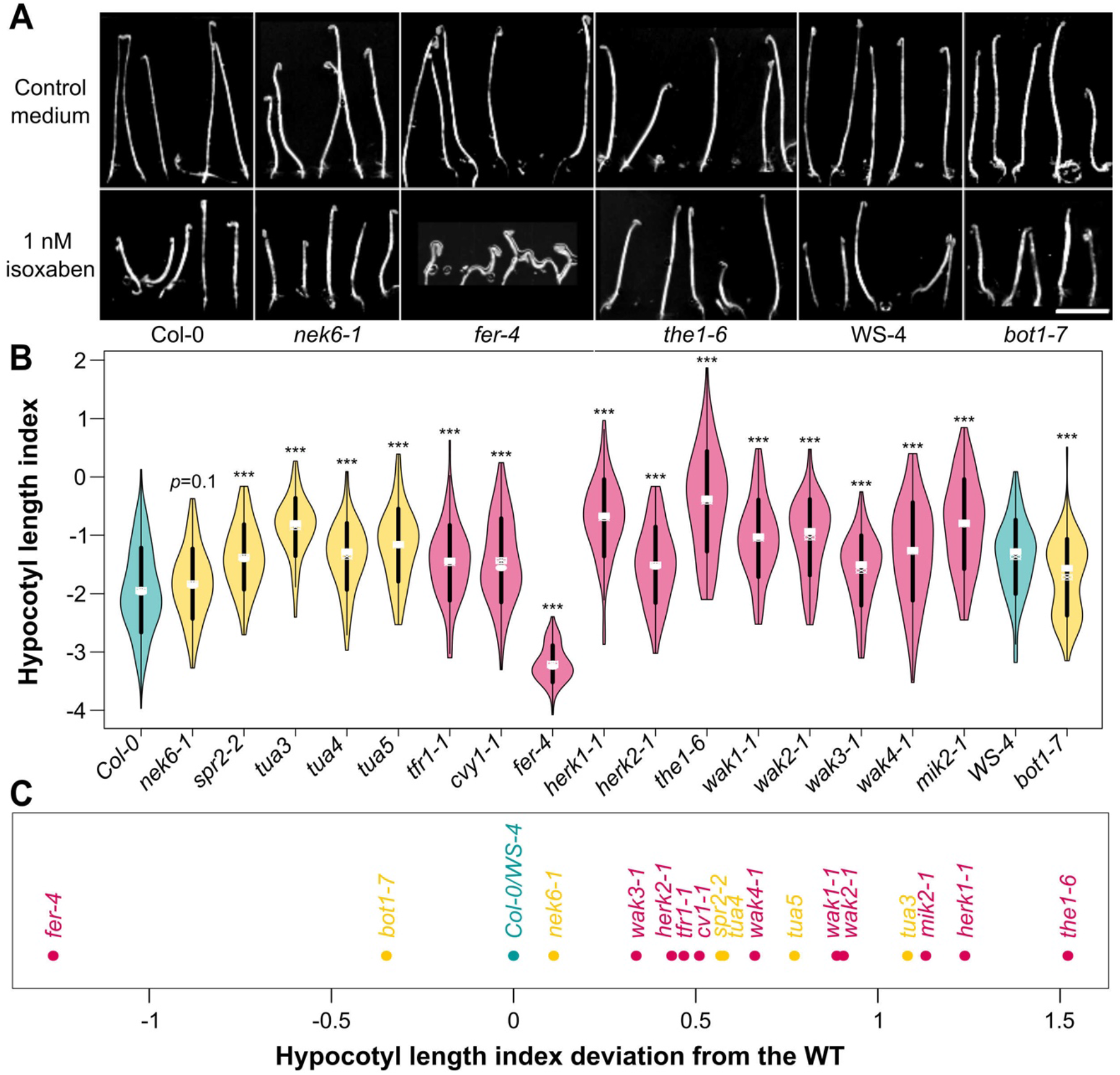
The impact of isoxaben on hypocotyl length reveals a differential contribution of receptor-like kinases in growth. **A.** Representative images of *Col-0*, *WS-4*, *fer-4*, *the1-6* and *bot1-7* etiolated seedlings grown with or without 1 nM isoxaben. Scale=1 cm. **B.** Hypocotyl length index (violin plot): distribution of isoxaben-grown hypocotyl length, normalized relative to the DMSO-grown ones. *p*-values (*p*) of Wilcoxon-Mann-Whitney test for the WT (in blue, Col-0, WS-4), the microtubule regulator (in orange, *nek6-1, spr2-2, tua3, tua4, tua5, bot1-7*), and the receptor-like kinase mutants (in pink, *tfr1-1, cvy1-1, fer-4, herk1-1, herk2-1, the1-4, the1-6, wak1-1, wak2-1, wak3-1, wak4-1, mik2-1*). **C.** Deviation of hypocotyl length index. The WT accessions (*Col-0* and *WS-4*) are labelled in blue. The microtubule regulator mutant lines (*nek6-1, spr2-2, tua3, tua4, tua5, bot1-7*) are labelled in orange. The receptor-like kinase mutants (*tfr1-1, cvy1-1, fer-4, herk1-1, herk2-1, the1-4, the1-6, wak1-1, wak2-1, wak3-1, wak4-1, mik2-1*) are labelled in pink.

10 out of the 11 receptor-like kinase mutants studied were less sensitive than the WT (*tfr1-1*, *cvy1-1*, *herk1-1*, *herk2-1, the1-6, wak1-1*, *wak2-1*, *wak3-1*, *wak4-1, mik2-1*; *p_tfr1-1_* <10^−3^; *p_cvy1-1_*<10^−3^; *p_herk1-1_*<10^−3^; *p_herk2-1_*<10^−3^; *p_the1-6_*<10^−3^; *p_wak1-1_*<10^−3^; *p_wak2-1_*<10^−3^; *p_wak3-1_*<10^−3^; *p_wak3-1_*<10_−3_; *p_mik2-1_*<10^−3^; n_*tfr1-1* DMSO_=80; n_*tfr1-1* iso_=83; n_*cvy1-1* DMSO_=109; n_*cvy1-1* iso_=115; n_*herk1-1* DMSO_=82; n_*herk1-1* iso_=70; n_*herk2-1* iso_=88; n_*herk2-1* iso_=106; n_*the1-6* DMSO_=57; n_*the1-6* iso_=91; n_*wak1-1* DMSO_=116; n_*wak1-1* iso_=89; n_*wak2-1* DMSO_=63; n_*wak2-1* iso_=78; n_*wak3-1* DMSO_=99; n_*wak3-1* iso_=89; n_*wak4-1* DMSO_=103; n_*wak4-1* iso_=86; n_*mik2-1* DMSO_=84; n_*mik2-1* iso_=96; n_Col-0 DMSO_=287; n_Col-0 iso_=271), whereas *fer-4* was significantly more sensitive (*p*_fer-4_<10^−3^; n_*fer-4* DMSO_=104; n_*fer-4* iso_=90) than the WT (Figure 2B). When plotting the deviation of each mutant from the WT phenotype, it appeared that among all the receptor-like kinase tested, *fer-4* and *the1-6* were the most affected mutants in their response to isoxaben, albeit in opposite trend: *fer-4* hypocotyl were more sensitive to isoxaben whereas *the1-*6 were less sensitive to isoxaben (Figure 2C). These results indicate that all the receptor-like kinases tested might be involved in the cellular response to a mechanical stress, with *FER* and *THE1* having the most clear-cut, and opposing, response.

To check whether these defects could be related to the microtubule response to stress, we performed the same analysis on microtubule regulator mutants. *bot1-7* (in *WS-4* ecotype) was more sensitive to the isoxaben treatment than the WT (*p_bot1-7_*<10^−3^; n_*bot1-7* DMSO_=99; n_*bot1-7* iso_=96; n_WS-4 DMSO_=101; n_WS-4 iso_=96) and thus fell in the same cluster as *fer-4.* The *nek6-1* mutant exhibited the same isoxaben sensitivity as the WT (*p_nek6-1_*=0.15; n_*nek6-1* DMSO_=91; n_*nek6-1* iso_=104; n_Col-0 DMSO_=287; n_Col-0 iso_=271)(Figure 2B). The *spr2-2* mutant was significantly less sensitive than the WT (p_*spr2-2*_<10^−3^; n_*spr2-2* DMSO_=99; n_*spr2-2* iso_=85; n_Col-0 DMSO_=287; n_Col-0 iso_=271) and thus fell in the same cluster as *the1-6.* Last, *tua3*, *tua4* and *tua5* were significantly less sensitive than the WT (*p_tua3_*<10^−3^; *p_tua4_*<10^−3^; *p_tua5_*<10^−3^; n_*tua3* DMSO_=93; n_*tua3* iso_=90; n_*tua4* DMSO_=83; n_*tua4* iso_=108; n_*tua5* DMSO_=72; n_*tua5* iso_=66; n_Col-0 DMSO_=287; n_Col-0 iso_=271).

Altogether, both pavement cell shape analysis and isoxaben test single FER out. Our data are also consistent with a scenario in which FER and katanin could belong to the same pathway. Yet, the isoxaben test reveals that the relation between wall defects and cortical microtubule response is not simple. In the following we decided to focus on FER and disentangle these apparent contradictions.

### Rescue test: Decreasing growth medium matrix potential rescues the burst cell phenotype in *fer-4*

It is commonly believed that WT seedlings display a shorter hypocotyl on isoxaben because cell wall defects are perceived and compensated through wall reinforcement, ultimately leading to reduced growth. Both wall reinforcement and reduced growth would prevent the cells from bursting. This notably builds on the analysis of *the1* mutant, which exhibits a longer hypocotyl than the WT on isoxaben because of the lack of wall sensing (Hématy et al., 2007). In *fer*, the hypocotyl is in contrast even shorter than the WT. Thus, either wall sensing is enhanced in *fer*, thus strongly repressing growth, or, in contrast, wall sensing is impaired in *fer*, even more than in *the1*, and cells burst before walls can even be reinforced. When looking closely at *fer-4* hypocotyls grown on isoxaben and stained with propidium iodide (PI), we observed many dead cells, as revealed by PI staining (Figure 3A). We calculated a bursting index, i.e. the percentage of burst cell area over total area of a field of epidermal cells in a given image and found it to be much higher in *fer-4* isoxaben-grown hypocotyls than in the WT (*p*<10^−3^; n_Col-0,isoxaben,0.7%agar_=16; n_*fer-4*,isoxaben,0.7%agar_=16, Figure 3B). This observation thus seems consistent with the latter scenario.

**Figure 3.**
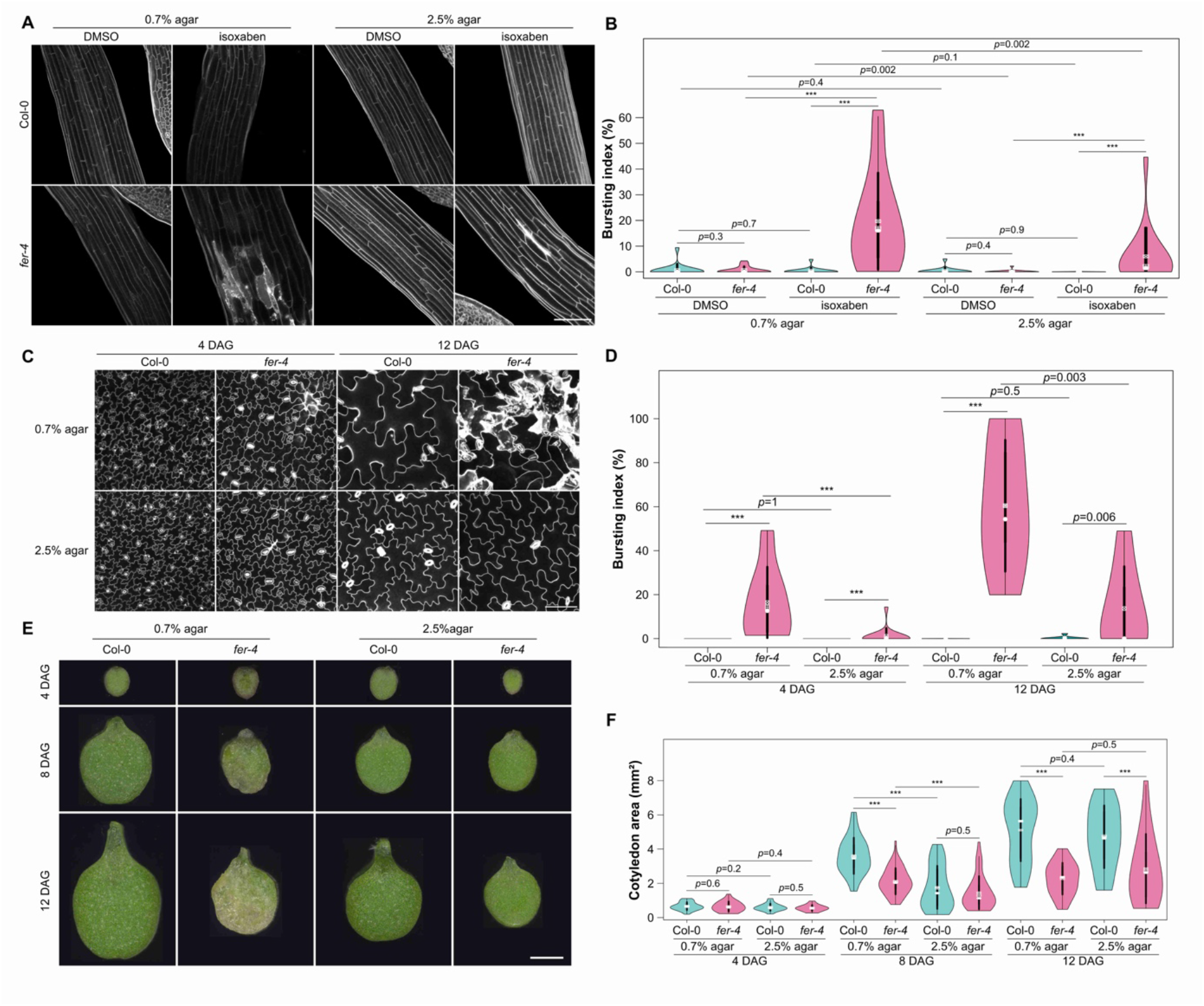
The *fer* phenotype can be partially rescued on 2.5% agar **A.** Representative confocal images of Col-0 and *fer-4* etiolated hypocotyls, from seedlings grown on a medium containing 1nM isoxaben with 0.7% or 2.5% agar (propidium iodide staining). Scale=100μm. **B.** Bursting index (violin plot) and *p*-values (*p*) of Wilcoxon-Mann-Whitney tests in *Col-0* and *fer-4* etiolated hypocotyls grown on a medium containing 1nM isoxaben with 0.7% or 2.5% agar. **C.** Representative confocal images of *Col-0* and *fer-4* pavement cells, from seedlings grown on a medium containing 0.7% or 2.5% agar at t=4 DAG and t=12 DAG (propidium iodide staining). Scale=100μm. **D.** Bursting index (violin plot) and *p*-values (*p*) of Wilcoxon-Mann-Whitney tests in *Col-0* and *fer-4* pavement cells, from seedlings grown on a medium containing 0.7% or 2.5% agar at t=4 DAG and t=12 DAG. **E.** Representative images of *Col-0* and *fer-4* cotyledons, grown on a medium containing 0.7% or 2.5% agar at t=4 DAG, t=8 DAG and t=12 DAG. Scale =1mm. **F.** Cotyledon area (violin plot) and *p*-values (*p*) of Wilcoxon-Mann-Whitney tests in *Col-0* and *fer-4* seedlings, grown on a medium containing 0.35%, 0.7%, 1.25% or 2.5% agar at t=4 DAG, t=8 DAG and t=12 DAG.

To test this hypothesis further, we reasoned that lowering the water potential in the medium should reduce water intake for seedlings, reduce tensile stress level (Verger et al., 2018), and in the end, reduce the number of burst cells. We thus tested different agar concentration in the medium and analysed the impact on *fer-4* phenotype. When increasing the agar concentration to 2.5%, the bursting index in hypocotyls was reduced by 70% in isoxaben-grown seedlings, supporting our hypothesis (p=0.002; n_fer-4,isoxaben,0.7%agar_=16; n_fer-4,isoxaben,2.5%agar_=16, Figure 3A and 3B).

We then took a closer look at the pavement cells of *fer-4* cotyledons grown in a medium containing a standard concentration of agar (0.7%) and over time. Many burst cells were present here as well, as marked by a strong PI coloration (Figure 3C). The bursting index increased over time, from 16±16% at 4 DAG (days after germination) to 60±30% at 12 DAG. At both time points, it was significantly superior to that of the WT (*p*_4DAG,0.7%agar_<10^−3^; *p*_12DAG,0.7%agar_<10^−3^; n_Col-0,4DAG,0.7%agar_=14, n_Col-0,12DAG,0.7%agar_=9, n_*fer-4*,4DAG,0.7%agar_ =14, n_*fer-4*,12DAG,0.7%agar_ 8; Figure 3D). When we increased the agar concentration to 2.5%, the bursting index in *fer-4* was still superior to that of the WT (*p*_4DAG,2.5%agar_<10^−3^; *p*_12DAG,2.5%agar_=0.063; n_Col-0,4DAG,2.5%agar_=15, n_Col-0,12DAG,2.5%agar_ 9, n_*fer-4*,4DAG,2.5%agar_ =15, n_*fer-4*,12DAG,2.5%agar_ =7; Figure 3D). However, the bursting index was dramatically reduced to 1±4% at 4 DAG (*p_fer-4_,*_4DA*G*_<10^−3^) and to 14±19% at 12 DAG (*p_fer-4_*_,12DAG_=0.003, Figure 3D). Once again, lowering the water availability for cells partially rescued the bursting cell phenotype in *fer-4*.

To check whether the apparent rescue could have a more global effect on organ shape, we measured *fer-4* cotyledon area over time on seedlings grown on media containing different agar concentrations (0.7%, 2.5%, Figure 3E and 3F). At a young stage (four days of light), *fer-4* cotyledons had comparable area as WT ones for every agar concentration (*p*_4DAG,0.7%agar_=0.63; *p*_4DAG,2.5%agar_=0.44; n_Col-0,4DAG,0.7%agar_=38; n_*fer-4*,4DAG,0.7%agar_=23; n_Col-0,4DAG,2.5%agar_=35; n_*fer-4*,4DAG,2.5%agar_=27, Figure 3E and 3F). After eight days of light, *fer-4* cotyledons were 35% smaller than the WT for seedling grown on a medium containing 0.7% agar (*p*_8DAG,0.7%agar_<10^−^ 3; n_Col-0,8DAG,0.7%agar_=45; n_*fer-4*,8DAG,0.7%agar_=56). Strikingly, *fer-4* cotyledons reached the same size as the WT for seedlings grown on 2.5% agar (*p*_8DAG,2.5%agar_=0.51; n_Col-0,8DAG,2.5%agar_=55; n_*fer-4*,8DAG,2.5%agar_=40). However, this trend was not maintained after twelve days of light (Figure 3E and 3F).

Thus, we find a correlation between the mechanical status of the *fer-4* cotyledons (as monitored by the frequency of cell burst) and growth. Taken together, these results suggest that the *fer-4* mutant is unable to sense the mechanical status of the tissue, and that the *fer-4* phenotype reflects a passive turgor-dependent wall defect. In other words, the *fer-4* mutation turns the plant from an active to a more passive material, where increased tensile stress leads to wall failure without feedback.

### Cortical microtubule orient with predicted tensile stress after ablation in *fer-4*

So far, all the tests suggest that FER is a major player in mechanosensing at the shoot, consistent with data obtained in the root (Shih 2014). However, it is not clear whether this involves the microtubule response to mechanical stress. To formally check this, we analysed the behaviour of cortical microtubules in response to local ablation in *fer-4*. From a mechanical standpoint, the sudden drop of turgor pressure in the ablated cells, together with epidermal tension (Kutschera and Niklas, 2007), generates a circumferential tensile stress around the dead cells (Hamant et al., 2008). Such a perturbation causes the cortical microtubules to reorient in the new maximal tensile stress direction (circumferential) around the ablation (Hamant et al., 2008)(Sampathkumar et al., 2014). We used the microtubule reporter *pPDF1∷mCit-MBD* to monitor the microtubule response in *Col-0* and *fer-4* cotyledons. As a 0.7% agar medium triggers widespread cell death in *fer-4* (Figure 3C and Supplementary Figure 3A-D), all tests were performed on 2.5% agar. We measured both the anisotropy and the average orientation of cortical microtubule arrays (relative to the ablation site) in cells surrounding the ablation (Figure 4 and Supplementary Figure 4).

**Figure 4.**
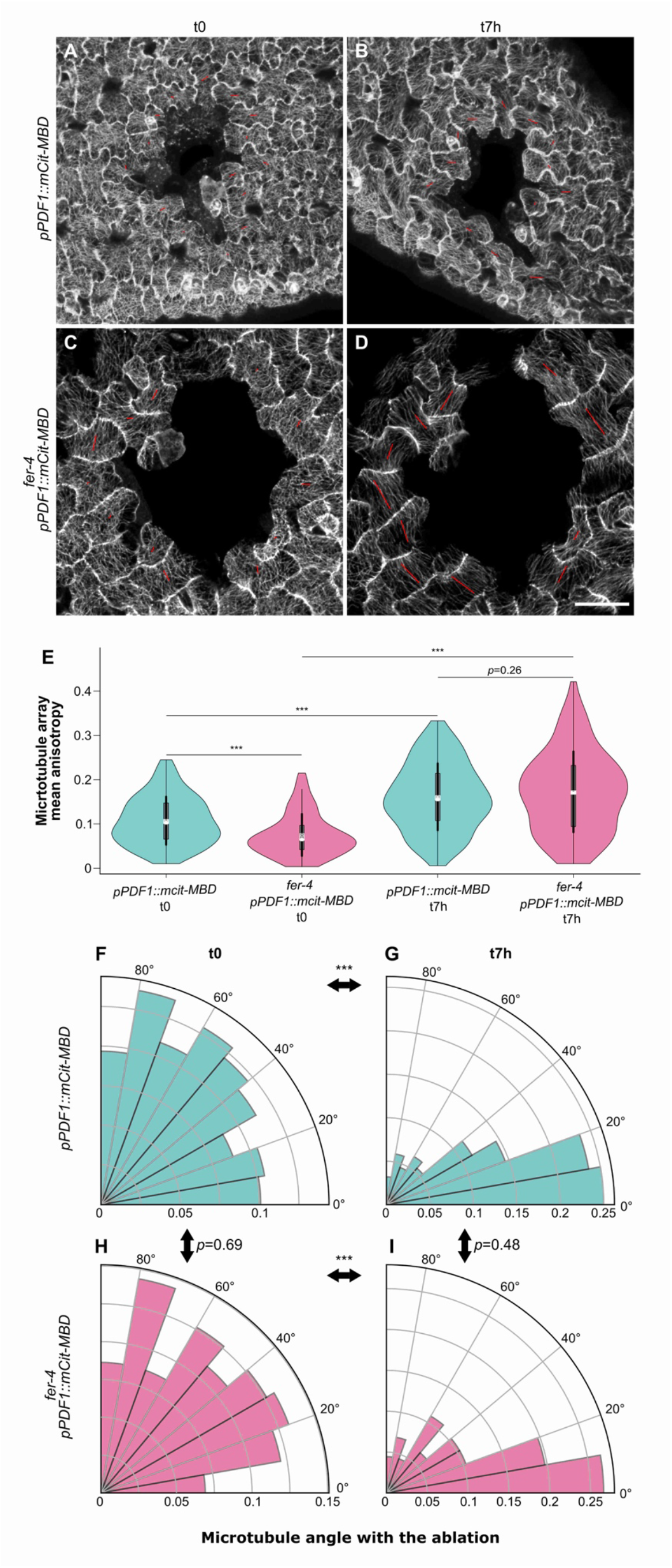
Cortical microtubule alignment with predicted maximal tensile stress in *fer* after ablation All seedlings were grown on 2.5% agar. **A-D.** Representative confocal images of *pPDF1∷mCit-MBD* (A, B) and *fer-4 pPDF1∷mCit-MBD* (C, D) pavement cells, immediately after an ablation (t0, A, C) and 7 hours later (t7h, B, D). The red bars indicate the average orientations of cortical microtubule arrays, and the length of the red bars is proportional to the anisotropy of the cortical microtubule arrays (using ImageJ FibrilTool). Scale=50μm. **E.** Anisotropy (violin plot) of cortical microtubule arrays and *p*-values (*p*) of Wilcoxon-Mann-Whitney tests in cells surrounding the ablation site in *pPDF1∷mCit-MBD* and *fer-4 pPDF1∷mCit-MBD* pavement cells, immediately after an ablation (t0) and 7 hours later (t7h). **F-I.** Cortical microtubule orientations (polar plots) and *p*-values (*p*) of Wilcoxon-Mann-Whitney tests in cells surrounding the ablation site in *pPDF1∷mCit*-*MBD* (F, G) and *fer-4 pPDF1∷mCit-MBD* (H,I) pavement cells, immediately after an ablation (t0, F, H)) and seven hours later (t7h, G, I).

The anisotropy (*a*) of cortical microtubule arrays was low in the WT immediately after ablation (*a_Col-0_*_,t0_=0.11; n_Col-0,t0_=248). It increased by 50 % seven hours later, consistent with the co-alignment of cortical microtubules with tensile stress (*a_Col-0_*_,t7h_=0.16; *p*<10^−3^; n_*Col-0*,t7h_=220; Figure 4E). In *fer-4*, cortical microtubule arrays were slightly less anisotropic (by 30%) than the WT at t=0h (*a_fer-4_,*_t0_=0.073; *p*<10^−3^; n_*fer-4*,t0_=174). After seven hours, the anisotropy of cortical microtubule arrays increased by 129 % and became comparable to that of the WT (*a_fer-4_,*_t7h_=0.17; *p*=0.26; *n_fer-4_,*_t7h_ =203, Figure 4E).

Immediately after the ablation, cortical microtubule arrays exhibited no preferred orientation (*o*) around the ablation, with an average angle of 47±25° for the WT, which was not significantly different from that of *fer-4* (*o_fer-4_,*_t0_=46±25°; *p*=0.69; Figure 4F and 4H, Supplementary Figure 4B). At t0, both distributions followed a uniform law on 0-90° (*p*_Col-0,t0_=0.5; *p_fer-4_*_,t0_=0.7; Kolmogorov-Smirnov test for uniformity). After seven hours, cortical microtubules did not follow a uniform distribution anymore (*p*_Col-0,t7h_<10^−3^; *p_fer-4_*_,t7h_<10^−3^; Kolmogorov-Smirnov test for uniformity) and became more circumferential around the ablation in both *Col-0* and *fer-4*, with no significant difference between the genotypes (*o_Col-0_*_,t7h_ =28±23°, *o_fer-4_,*_t7h_=31±25°; *p*=0.48, Figure 4G and 4I, Supplementary Figure 4).

Similar trends could be observed when using the *p35S∷GFP-TUB* microtubule marker line: cortical microtubule orientation appeared circumferential around ablations in both WT and *fer-4* (Supplementary Figure 5). However, the diffuse fluorescent signal hindered quantitative analysis with FibrilTool. Because the microtubule response to ablation is comparable in *fer-4* and in the WT, this formally shows that the microtubule response to stress can be independent from FER.

### FERONIA and microtubules independently contribute to the response to mechanical stress

Our ablation results may seem at odds with the fact that *fer-4* and *bot1-7* belong to the same cluster when analysing pavement cell circularity (see Figure 1D-E). We thus revisited our quantification of pavement cell shape to identify other shape descriptors amenable to discriminate *bot1-7* and *fer-4*. We focused on lobe size in pavement cells. In the katanin mutant *bot1-7*, which displayed a higher circularity than the WT (see Figure 1), the average basal lobe width was 3% smaller than that of the WT (*p*<10^−3^; *n*_WS-4_=294; *n_bot1-7_*=252; Figure 5A). Although this is a rather small difference, this is consistent with a reduced microtubule response to stress in the katanin mutant. In contrast, the average basal lobe width in *fer-4* was 12% larger than that of the WT (*p*<10^−3^; *n*_Col-0_=428; *n_fer-4_*=321; Figure 5A). Similar trends in basal lobe width were observed when seedlings were grown on 2.5% agar, albeit with lower values, also consistent with a reduced microtubule response to mechanical stress in such conditions (Supplementary Figure 6). Thus, both *fer-4* and *bot1-7* pavement cells exhibit higher circularity than the WT through different geometries. Pavement cell shape may thus reflect different responses to stress: reduced microtubule dynamics in *bot1-7* would generate smaller lobes, whereas weaker walls in *fer-4* would increase stress levels, leading to hyper-aligned cortical microtubules and larger lobes. Such hyper-aligned cortical microtubules can be observed in *fer-4* pavement cells when seedlings are grown on 0.7% agar (see Supplementary Figure 3C). Alternatively, and non-exclusively, the presence of dead cells in *fer-4* may affect the stress pattern, and thus the cortical microtubule response, further increasing the circularity of pavement cells.

**Figure 5.**
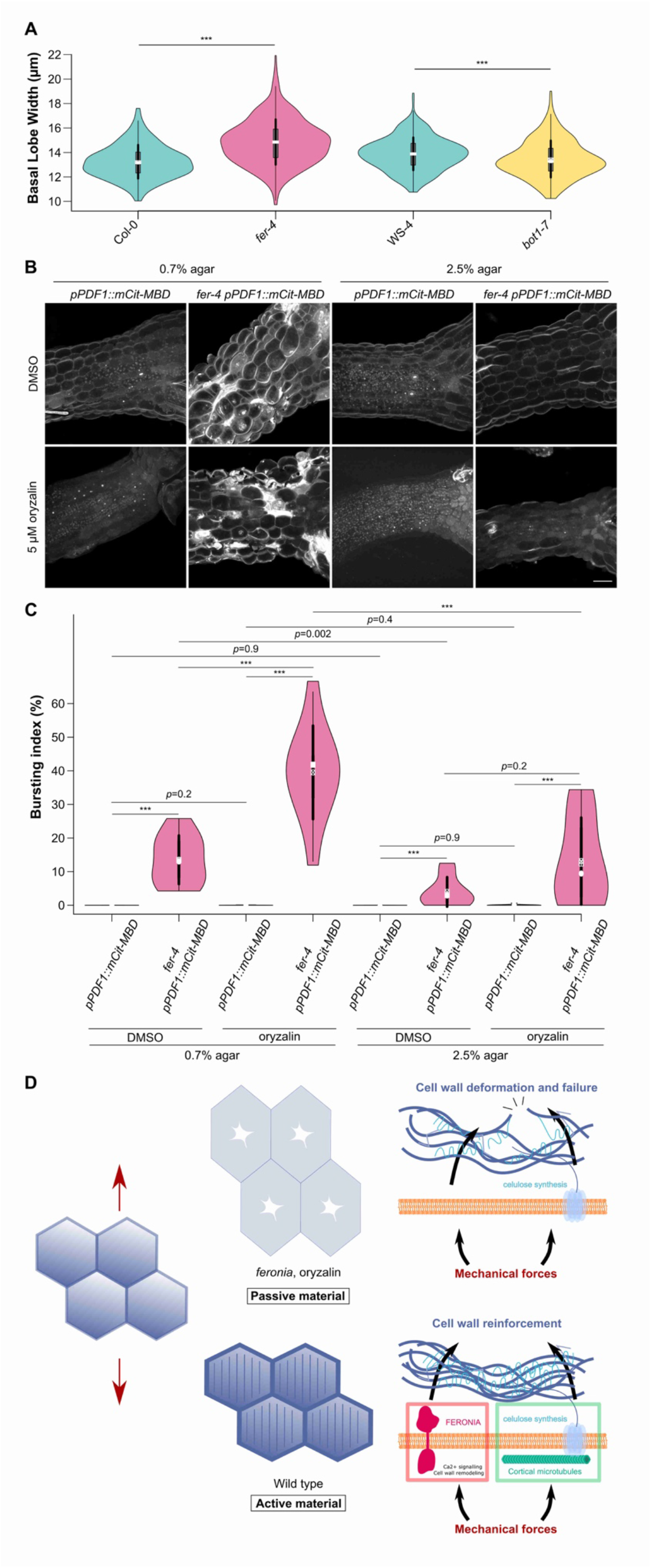
FER and microtubules independently contribute to the response to mechanical stress **A.** Basal lobe width (violin plot) of pavement cells and *p*-values (*p*) of Dunn tests for the WT (*Col-0, WS-4*), katanin mutant (*bot1-7*), and *fer-4*. Seedlings were grown on 0.8% agar. **B.** Representative confocal images of *pPDF1∷mCit-MBD* and *fer-4 pPDF1∷mCit-MBD* hypocotyls grown on 0.7% and 2.5% agar with and without 5 μm of oryzalin. Scale bar=50μm. **C.** Bursting index (violin plot) and *p*-values (*p*) of Wilcoxon-Mann-Whitney tests in *pPDF1∷mCit-MBD* and *fer-4 pPDF1∷mCit-MBD* hypocotyls grown on 0.7% or 2.5% agar, with and without 5 μm of oryzalin. **D.** In the WT (bottom), cells resist mechanical stress (red arrows) by two independent pathways (microtubule-dependent cellulose synthesis and FER-dependent wall reinforcement). In absence of both FER and microtubules (top), cells deform like passive materials and ultimately burst.

Treatment with the microtubule depolymerizing drug oryzalin has previously been shown to induce cell bursting events in the largest cells at the shoot apical meristem (Sapala et al., 2018), mimicking the bursting cell phenotype we observe in hypocotyls and cotyledons in *fer-4*. If both pathways are truly independent, then one should expect additive behaviors. To test this prediction, we observed hypocotyl growth when both pathways are down, by applying oryzalin on *fer-4* mutants. On 0.7% agar, bursting cells appeared in both oryzalin-treated hypocotyls and control ones in *fer-4.* The bursting index increased by 80% in oryzalin-treated *fer-4* hypocotyls, consistent with an additive role of both pathways in wall integrity (*p_fer-4_,*_0.7%agar_=0.001; n_*fer-4*,DMSO,0.7%agar_=10; n_*fer-4*,oryzalin,0.7%agar_=11, Figure 5B-C). To relate this phenotype to stress levels, we performed the same experiment on 2.5% agar, and found a reduction in the bursting cells in all conditions (by 87% on DMSO; *p_fer-4_,*_DMSO_=0.002; n_*fer-4*,DMSO,0.7%agar_=10; n_*fer-4*,DMSO,2.5%agar_=9; by 80% on oryzalin; *p_fer-4_,*_oryzalin_<10^−3^; n_*fer-4*,oryzalin,0.7%agar_=11; n_*fer-4*,oryzalin,2.5%agar_=10; Figure 5B-C). This further confirms that FERONIA and microtubules independently contribute to the response to stress.

## Discussion

From a reverse genetic screen, we show that FERONIA plays a primary role in the cell response to mechanical stress in the shoot. We also find that the microtubule response to stress does not depend on FER. This provides a scenario in which the mechanical feedback in the shoot involves two independent modules, cortical microtubules and FER (Figure 5D).

This result is consistent with recent findings showing how pectin and cellulose deposition are controlled with largely disconnected networks: Pectins are important for the initiation of pavement cell formation, and the deposition of cellulose microfibril rather amplifies a pre-existing mechanical pattern (Bidhendi et al., 2019). Similarly, mechanical polarities in growing hypocotyl cells precedes the cellulose-derived mechanical anisotropy in hypocotyl cells (Peaucelle et al., 2015).

We also provide evidence that the *fer* mutant can be largely rescued though changes in the mechanical environment of the plant despite its central and somewhat pleiotropic role in plant development. This echoes recent findings where essential regulators are found optional, when challenged with different mechanical environments. This is the case for instance for key regulators of pectin synthesis, where decreasing tensile stress can rescue cell-cell adhesion defects in the *quasimodo* mutants (Verger et al., 2018). This was also shown for the katanin mutant, where increasing tensile stress levels with isoxaben treatment can generate WT-like cortical microtubule arrays (Uyttewaal et al., 2012). More recently, this was even extended to signaling in the context of apical hook formation: *arf7 arf19* auxin transduction mutant exhibit a WT phenotype when mechanically constrained (Baral et al., 2020).

If the microtubule response to stress does not depend on FER, what could be the most relevant mechanotransduction pathway? As shown with optical tweezers, pulling on microtubules promote their polymerization *in vitro* (Franck et al., 2007)(Trushko et al., 2013). Cortical microtubules also align with predicted maximal tension when protoplasts are confined in rectangular microwells and pressurized by hypo-osmotic conditions (Colin et al., 2020). These data, together with the fer-independent microtubule response to stress, further support the hypothesis that microtubules may be their own mechanosensor (Hamant et al., 2019).

Conversely, the final microtubule (and cellulose microfibril) alignment likely results from a combination of several cues, beyond tensile stress. Cell geometry can affect microtubule behaviour, independent of cortical tension. In particular, due to their high persistence length, microtubules tend to become longitudinal *in vitro* (Cosentino Lagomarsino et al., 2007) or in depressurized protoplasts (Durand-Smet et al., 2020) (Colin et al., 2020). Furthermore, cell edge factors can affect cortical microtubule behaviour, leading to cell-scale aligned arrays (Ambrose et al., 2011)(Chakrabortty et al., 2018)(Kirchhelle et al., 2019). Although microtubule, FER, stress and cell geometry can be uncoupled in experiments, their interplay may provide synergies *in vivo*. In particular, the deposition of matrix material in the wall depends on exocytosis, which is also promoted by membrane tension. Conversely, affecting matrix deposition may weaken the wall and increase the tensile stress levels. Consistently, microtubule arrays exhibit an enhanced anisotropy when the cell edge GTPase Rab-A5c dependent trafficking is affected in roots (Kirchhelle et al., 2019).

All living organisms constantly sense and respond to mechanical stress. By impairing two major mechanosensing players, we show what plants looks like when cells become unable to respond to stress. We find that cells switch to a more passive mode, like a balloon or a soap bubble: cell behaviour scales with stress properties, and usually they die ultimately. Defining living systems as active material is thus particularly suited to understand how cells and tissue behave in response to fluctuations in their physical environment. The physics of active matter, and the associated mechano-chemical couplings, may be essential to revisit the role of many plant signaling pathways too.

## Material and Methods

### Plant material and growth conditions

All plants were in the Columbia-0 (Col-0) ecotype, except the *bot1-7* mutant, which was in the Wassilewskija (WS-4) ecotype (Supplementary Table 1).

Seeds were surface sterilized and individually sown on Murashige and Skoog medium (MS medium, Duchefa, Haarlem, the Netherlands) or Arabidopsis medium (custom-made Duchefa ‘Arabidopsis’ medium (DU0742.0025, Duchefa Biochemie) with different agar concentrations (see Supplementary Table 2 for detailed description of the different medium used). For drug treatments, mediums were supplemented with isoxaben (Sigma) or oryzalin (Chem Service Inc.) from stock solutions in dimethyl sulfoxide (DMSO, Sigma). All plants were placed 2 days in the dark at 4°C then transferred in a 20°C long-days growth chamber. When seedlings were maintained in the dark, petri dishes were covered with aluminium foil in the 20°C long-days growth chamber.

### Image acquisition

Samples were imaged with either a SP8 confocal microscope (Leica Microsystems) equipped with a 25x long-distance water objective (NA = 0.95), an Epson Perfection 2400 scanner, or a Leica MZ12 microscope (As specified in Supplementary Table 2). Samples were stained for 10 minutes with a propidium iodide solution (Sigma; PI stains wall pectins and thus marks cell contours). Ablations (Figure 4) were performed manually with a fine needle (Minutien pin, 0.15 mm rod diameter, 0.02 mm tip width, RS-6083–15, Roboz Surgical Instrument Co.) as described in (Verger et al., 2018). In all confocal microscopy images, 0.5 μm thick optical slices were acquired.

For every experiment, three biological replicates or more were obtained. Col-0 or WS-4 seedlings were included as controls in all experiments and replicates. Note that all of the Col-0 images used in the first figure of this study have also been used in (Erguvan et al., 2019), as templates to introduce the SurfCut ImageJ tool (see below in Image analysis) for cell contour extraction.

### Image analysis

Pavement cell shape were obtained by first processing confocal images with MorphoGraphX (http://www.mpipz.mpg.de/MorphoGraphX) (Barbier de Reuille et al., 2015) to obtain cell contours in a 2.5D epidermal surface (Figures 1 and 5A) or SurfCut (https://github.com/sverger/SurfCut) (Erguvan et al., 2019) to extract the flattened cell contours (Figure 3E). The cell contour images were then processed with PaCeQuant (Möller et al., 2017), an ImageJ plugin quantifying up to 27 shape descriptors of pavement cells. Hypocotyl length (Figure 2) were measured manually with ImageJ (https://fiji.sc/). Cell burst area (Figure 3 and 5C-D) was measured manually with ImageJ after extracting the flattened cell contours with SurfCut (https://github.com/sverger/SurfCut) (Erguvan et al., 2019). Cotyledon area (Figure 3) was measured manually with ImageJ. Microtubule organization (Figure 4) was quantified with FibrilTool (Boudaoud et al., 2014) after flattening the images with SurfCut and denoising them (ROF Denoise, Theta=25) in ImageJ, as performed in (Verger et al., 2018). Ablations (Figure 4) were performed manually with a fine needle (Minutien pin, 0.15 mm rod diameter, 0.02 mm tip width, RS-6083–15, Roboz Surgical Instrument Co.) as described in (Verger et al., 2018). After image analyses, the luminosity and contrast of all images presented in this study were enhanced to help visualization.

### Statistical analysis

Statistical analyses were performed with R software (https://www.R-project.org). The sample size is indicated in the main text and in the figure legends. For pavement cell shapes (Figures 1, 5A and 6A), we used PaCeQuantAna, the R script that accompanies the PaCeQuant analysis (Möller et al., 2017), and the FactoMineR and factoextra R libraries for the PCA (Lê et al., 2008). Violin plots were shown with the corresponding *p*-value of Kruskal-Wallis tests. For hypocotyl length (Figure 2), the control distribution of hypocotyl length was standardized and the same parameters (μ_ctrl_, ơ_ctrl_) used to shift the isoxaben distribution similarly (Sup. Figure 2). A Wilcoxon–Mann–Whitney test was then performed on the shifted distributions of the mutant and of the WT for each genotype. For the orientation of microtubules, a Kolmogorov-Smirnov test was performed to compare the angle distributions to a uniform distribution between 0 and 90°. All other quantitative measures were compared using Wilcoxon–Mann–Whitney tests. As a Wilcoxon-Mann-Whitney test can be directed or not, the p-value shown in all experiments with a Wilcoxon-Mann-Whitney test was that of a non-directed test for non-significant p-value (to ensure that neither distribution was higher than the other) and that of a directed test in the significant direction for a significant p-value. Differences with *p*-values that were under 1% were considered significant, and those between the commonly used thresholds of 5% and 1% were considered as tendencies. All ± values referred to the standard deviation of the distribution.

## Acknowledgments

We are thankful to our colleagues at the plant reproduction and development lab for their comments and feedback on this manuscript. We thank Platim for help with imaging. This work was supported by the European Research Council (ERC-2013-CoG-615739 ‘MechanoDevo’.

