## Supplementary information for "Turning plants from passive to active material: FERONIA and microtubules independently contribute to mechanical feedback"

### Supplementary Table 1. Accessions

| ID | Line | Full gene name | AGI | Ontology | Ecotype | Reference |
|---|---|---|---|---|---|---|
| *nek6-1* | SALK_152782 | *NIMA (NEVER IN MITOSIS, GENE A)-RELATED 6* | AT3G44200 | Tubulin kinase | Col-0 | Motose *et al.*, 2008. Plant J Cell Mol Biol 54(5):829–844. |
| *spr2-2* | EMS CS6549 | *SPIRAL2* | AT4T27060 | MAP | Col-0 | Shoji *et al.*, 2004. Plant Physiol 136(4):3933–3944. |
| *tua3(D205N)* | EMS CS68878 | *TUBULIN ALPHA-3* | AT5G19770 | Tubulin | Col-0 | Ishida et al., 2007. Proc Natl Acad Sci U S A 104(20):8544–8549. |
| *tua4(S178D)* | EMS CS68881 | *TUBULIN ALPHA-4* | AT1G04821 | Tubulin | Col-0 | Ishida et al., 2007. Proc Natl Acad Sci U S A 104(20):8544–8549. |
| *tua5(D251N)* | EMS CS68884 | *TUBULIN ALPHA-5* | AT5G19780 | Tubulin | Col-0 | Ishida et al., 2007. Proc Natl Acad Sci U S A 104(20):8544–8549.. |
| *tfr1-1* | GK649 – E11-67 | *THESEUS-FERONIA-RELATED1* | AT5G24010 | CrRLK | Col-0 | This study, provided by H. Höfte |
| *cvy1-1* | SALK_018797 | *CURVY1* | AT2G39360 | CrRLK | Col-0 | Gachomo *et al.*, 2014. BMC Plant Biol 14:221. |
| *fer-4* | GABI 106_A06 | *FERONIA* | AT3G51550 | CrRLK | Col-0 | Duan et al., 2010. Proc Natl Acad Sci U S A 107(41):17821–17826. |
| *herk1-1* | SALK_008043 | *HERCULES RECEPTOR KINASE 1* | AT3G46290 | CrRLK | Col-0 | Guo et al., 2009. Proc Natl Acad Sci U S A 106(18):7648–7653. |
| *herk2-1* | SALK_105055 | *HERCULES RECEPTOR KINASE 2* | AT1G30570 | CrRLK | Col-0 | Guo et al., 2009. Proc Natl Acad Sci U S A 106(18):7648–7653. |
| *the1-6* | EMS (sup ctl1-2) | *THESEUS1* | AT5G54380 | CrRLK | Col-0 | Merz *et al.*, 2017. J Exp Bot 68(16):4583–4593. |
| *wak1-1* | SALK_107175 | *CELL WALL-ASSOCIATED KINASE 1* | AT1G21250 | WAK | Col-0 | He *et al.*, 1996. J Biol Chem 271(33):19789–19793. |
| *wak2-1* | SAIL_286_E03 | *CELL WALL-ASSOCIATED KINASE 2* | AT1G21270 | WAK | Col-0 | He *et al.*, 1996. J Biol Chem 271(33):19789–19793. |

| | | | | | | |
|---|---|---|---|---|---|---|
| *wak3-1* | SALK_071999 | *CELL WALL-ASSOCIATED KINASE 3* | AT1G21240 | WAK | Col-0 | He et al., 1999. Plant Mol Biol 39(6):1189–1196. |
| *wak4-1* | SAIL_1156_F08 | *CELL WALL-ASSOCIATED KINASE 4* | AT1G21210 | WAK | Col-0 | He et al., 1999. Plant Mol Biol 39(6):1189–1196. |
| *mik2-1* | SALK_061769 | *MDIS1-INTERACTING RECEPTOR LIKE KINASE2* | AT4G08850 | LRR-RLK | Col-0 | Wang *et al.,* 2016. Nature 531(7593):241–244. |
| *bot1-7* | deletion of 19 bp | *KATANIN 1* | AT1G80350 | Katanin | WS-4 | Bichet *et al.,* 2001. Plant J 25(2):137–148. |
| *pPDF1::mCit-MBD* | Translational fusion protein | | | Microtubule marker | Col-0 | Armezzani *et al.,* 2018. Development 145. |
| *fer-4 pPDF1::mCit-MBD* | Translational fusion protein | | | Microtubule marker | Col-0 | Cross, this study |
| *p35S::GFP-TUB* | Translational fusion protein | | | Microtubule marker | Col-0 | Lin et al., 2018) BioRxiv. https://doi.org/10.1101/269647 |
| *fer-4 p35S::GFP-TUB* | Translational fusion protein | | | Microtubule marker | Col-0 | Lin et al., 2018) BioRxiv. https://doi.org/10.1101/269647 |

Abbreviations: MAP: Microtubule Associated Protein, CrRLK: Catharanthus Receptor-Like Kinase, WAK: Wall Associated Kinase, LRR-RLK: Leucin Rich Repeat Receptor-Like Kinase

**Supplementary Table 2.** Growth and imaging conditions

| Figure | Panel | Medium | Growth conditions | Imaging |
|---|---|---|---|---|
| Figure 1 | A-D | Medium A | Condition 1 | Imaging a |
| Figure 2 | A-C | Medium B | Condition 2 | Imaging b |
| Figure 3 | A-B | Medium C | Condition 2 | Imaging c |
| Figure 3 | C-D | Medium D | Condition 3 | Imaging a |
| Figure 3 | E-F | Medium D | Condition 3 | Imaging f |
| Figure 4 | A-I | Medium D | Condition 1 | Imaging d |
| Figure 5 | A | Medium A | Condition 1 | Imaging a |
| Figure 5 | B-C | Medium E | Condition 4 | Imaging e |

| **Growth medium** | |
|---|---|
| Medium A | MS medium with 0.8% agar, 1% sucrose, and no vitamin |
| Medium B | MS medium with 0.8% agar, no sucrose, and no vitamin supplemented with 0.1 nM of isoxaben diluted in DMSO, or the same volume of DMSO (control) |
| Medium C | MS medium with 0.8% or 2,5% agar, no sucrose, and no vitamin supplemented with 0.1 nM of isoxaben diluted in DMSO, or the same volume of DMSO |
| Medium D | Arabidopsis medium with 0,7% or 2,5% agar and no vitamin |
| Medium E | Arabidopsis medium with 0,7% or 2,5% agar, no vitamin, and supplemented with 5 µM of oryzalin diluted in DMSO, or the same volume of DMSO |

| **Growth conditions** | |
|---|---|
| Condition 1 | 8 hours light, 3 days darkness, 3 days light |
| Condition 2 | 8 hours light, 4 days darkness |
| Condition 3 | 4 days light or 12 days light |
| Condition 4 | 5 days light |

| **Imaging**<br>In all confocal microscopy image acquisition, optical sections were 0.5 µm thick. | |
|---|---|
| Imaging a | Samples were transferred to new medium plates, stained with a PI solution (dilution 1/10), and rinsed twice. Cotyledons were manually set in flat position with forceps under a binocular stereoscopic microscope and imaged with a long distance 25x objective (Leica SP8). |
| Imaging b | The petri dish lid was removed and the petri dish was scanned with an office scanner. |
| Imaging c | Samples were stained with PI (1/100 dilution), rinsed once, mounted in water between slides and coverslips, and imaged with a long distance 25x objective (Leica SP8). |
| Imaging d | Samples were transferred to new medium plates. Cotyledons were manually set in flat position with forceps under a binocular stereoscopic microscope, and imaged with a 25x long distance objective (Leica SP8). |
| Imaging e | Samples were stained with PI (1/100 dilution), rinsed once, mounted in medium and water between slides and coverslips, and imaged with a long distance 25x objective (Leica SP8). |
| Imaging f | Dissected cotyledons were mounted in water between slides and coverslips and imaged with a binocular stereomicroscope (Leica MZ12). |

**Supplementary Figure 1**

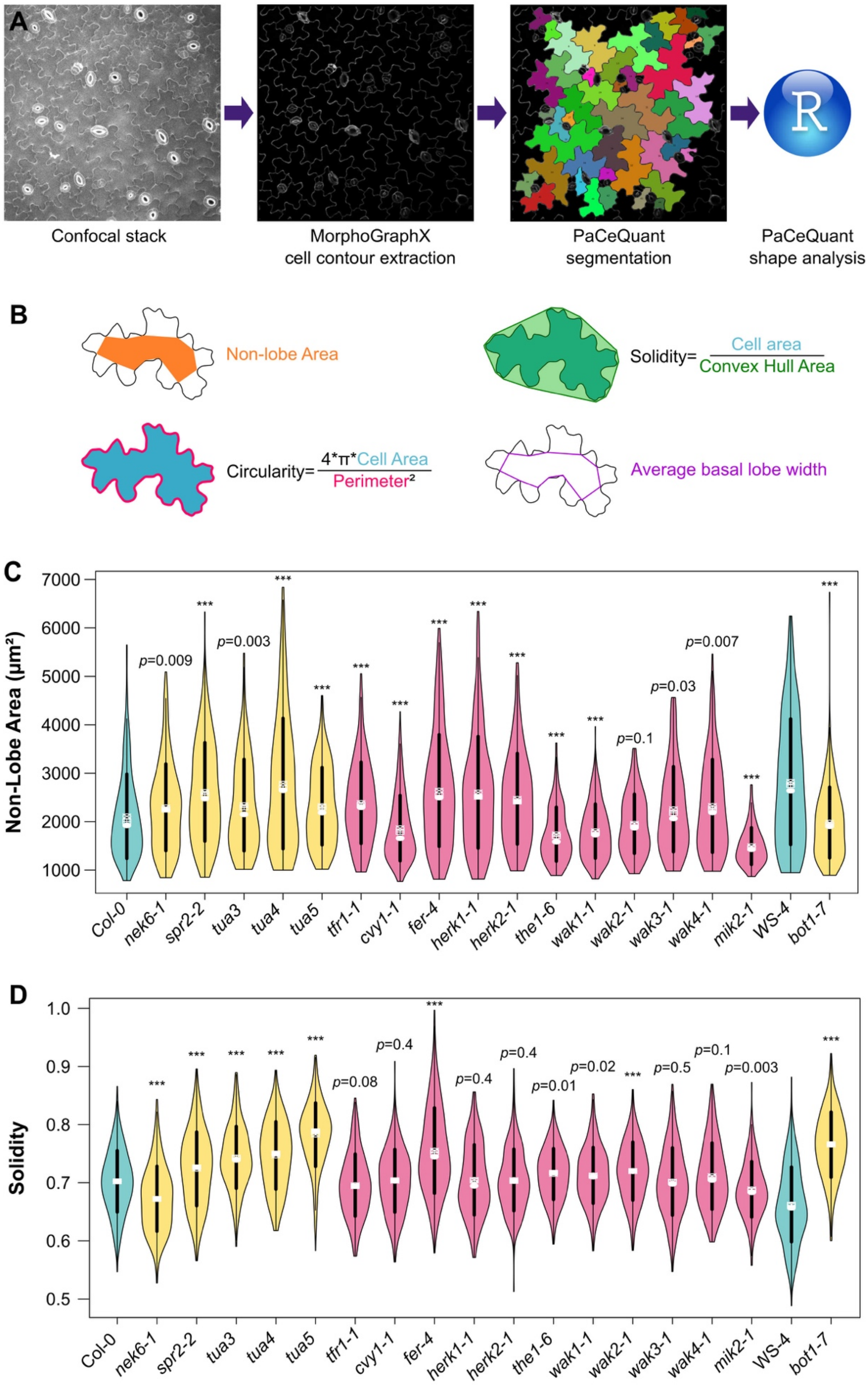

**Supplementary Figure 1.** Analysis of pavement cell shapes

A. Pipeline for the extraction and analysis of pavement cell shape. Epidermal signal was first extracted with MorphoGrahX, then segmented with the PaCeQuant ImageJ plugin, which extracted 27 pavement cell shape descriptors (parameters). Finally, the results were processed by PaCeQuantAna (the PaCeQuant R script).

B. Main shape descriptors of pavement cells. Non-lobe area (in $\mu m^2$) stands for the area of a cell without the lobes. Circularity is the area of a cell divided by its square perimeter, with a normalization to have a maximal circularity of 1 for a circle. Solidity is the area of a cell divided by the area of its convex hull (convex polygon with the smallest area including the whole cell). Average basal lobe width (µm) stands for the average length of the bases of a cell's lobes.

C. Non-lobe area (violin plots) of pavement cells and *p*-values (*p*) of Dunn tests in the WT (in blue, *Col-0, WS-4*), in microtubule associated mutants (in orange, *nek6-1, spr2-2, tua3, tua4, tua5, bot1-7*) and in receptor-like kinase mutants (in pink, *tfr1-1, cvy1-1, fer-4, herk1-1, herk2-1, the1-4, the1-6, wak1-1, wak2-1, wak3-1, wak4-1, mik2-1*).

D. Solidity (violin plots) of pavement cells and *p*-values (*p*) of Dunn tests in the WT (in blue, *Col-0, WS-4*), in microtubule associated mutants (in orange, *nek6-1, spr2-2, tua3, tua4, tua5, bot1-7*) and in receptor-like kinase mutants (in pink, *tfr1-1, cvy1-1, fer-4, herk1-1, herk2-1, the1-4, the1-6, wak1-1, wak2-1, wak3-1, wak4-1, mik2-1*).

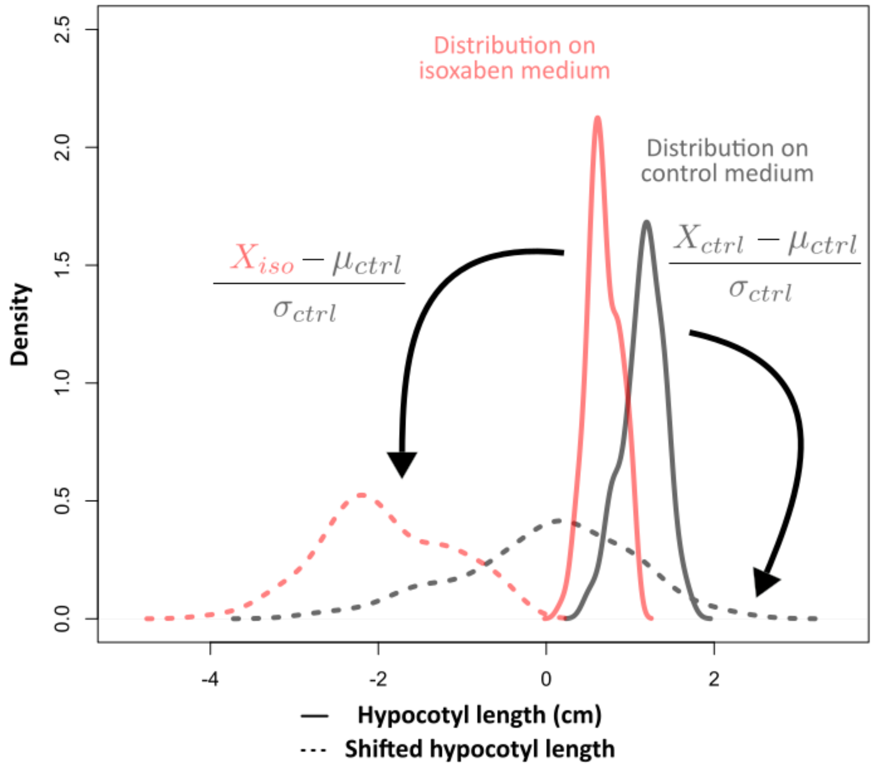


**Supplementary Figure 2.** Definition of the hypocotyl length index

The continuous black and red lines represent the distribution of the actual measurement of hypocotyl lengths for the treated (red) and control (black) samples. In this example, the values are between 0 and 2 cm and the average values for the treated sample are lower. However when comparing different genotypes, the average length of the untreated (control) mutant samples could already be lower or higher than the control WT samples thus hampering a direct comparison of the effect of the treatment on the mutant. In order to allow statistical comparison between the samples we normalized the distribution of hypocotyl lengths in the presence of isoxaben according to the distribution of hypocotyl lengths in control conditions, following the standard score method. To do so, the parameters of the control distribution (DMSO, mean: $\mu_{ctrl}$, standard deviation: $\sigma_{ctrl}$) are shifted around 0 (dashed black line) and are used to shift the isoxaben distribution to the same extent (dashed red line). With this method, the control samples for all genotypes have a value around 0, which then allows to reveal the effect of the treatment on the different mutants by comparing the means of the normalized treated samples (The hypocotyl length index; Figure 2B). To further ease the comparison, the differences between WT and mutants are then plotted as the Hypocotyl length index deviation from the WT, which are the values obtained when subtracting the mutant treated normalized mean to the WT treated normalized mean. This generates a negative value in the hypocotyl length index for mutants being more sensitive to the treatment and a positive value for mutants less sensitive to the treatment (Figure 2C).

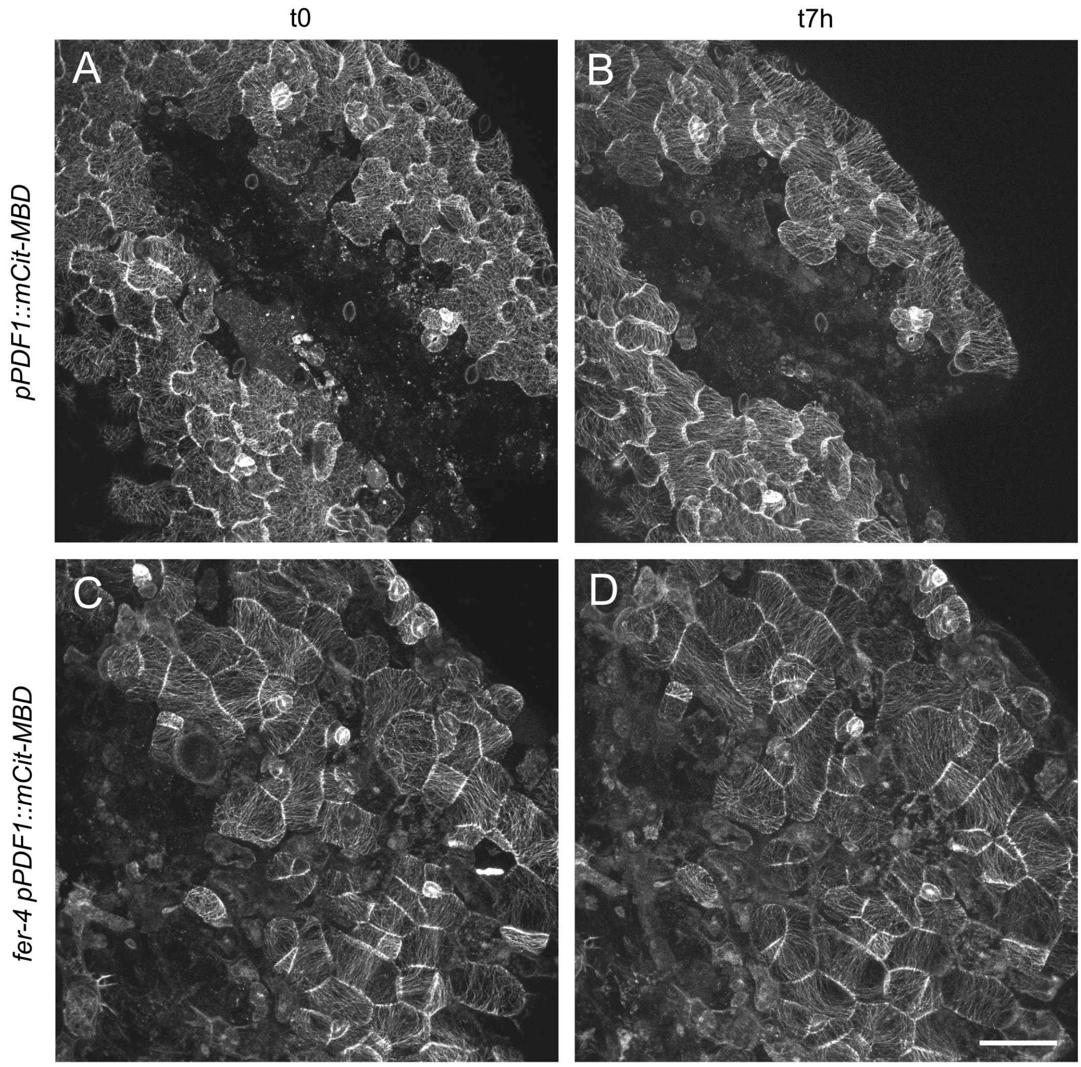


**Supplementary Figure 3.** Ablations on 0.7% agar medium

Representative confocal images of *pPDF1::mCit-MBD* (A, B) and *fer-4 pPDF1::mCit-MBD* (C, D) pavement cells from seedlings grown on a medium containing 0.7% agar, immediately after ablation (t0, A, C) and seven hours later (t7h, B, D). Note the presence of many dead cells and the strong alignment of cortical microtubules at t0 in *fer-4*. Scale=50µm.

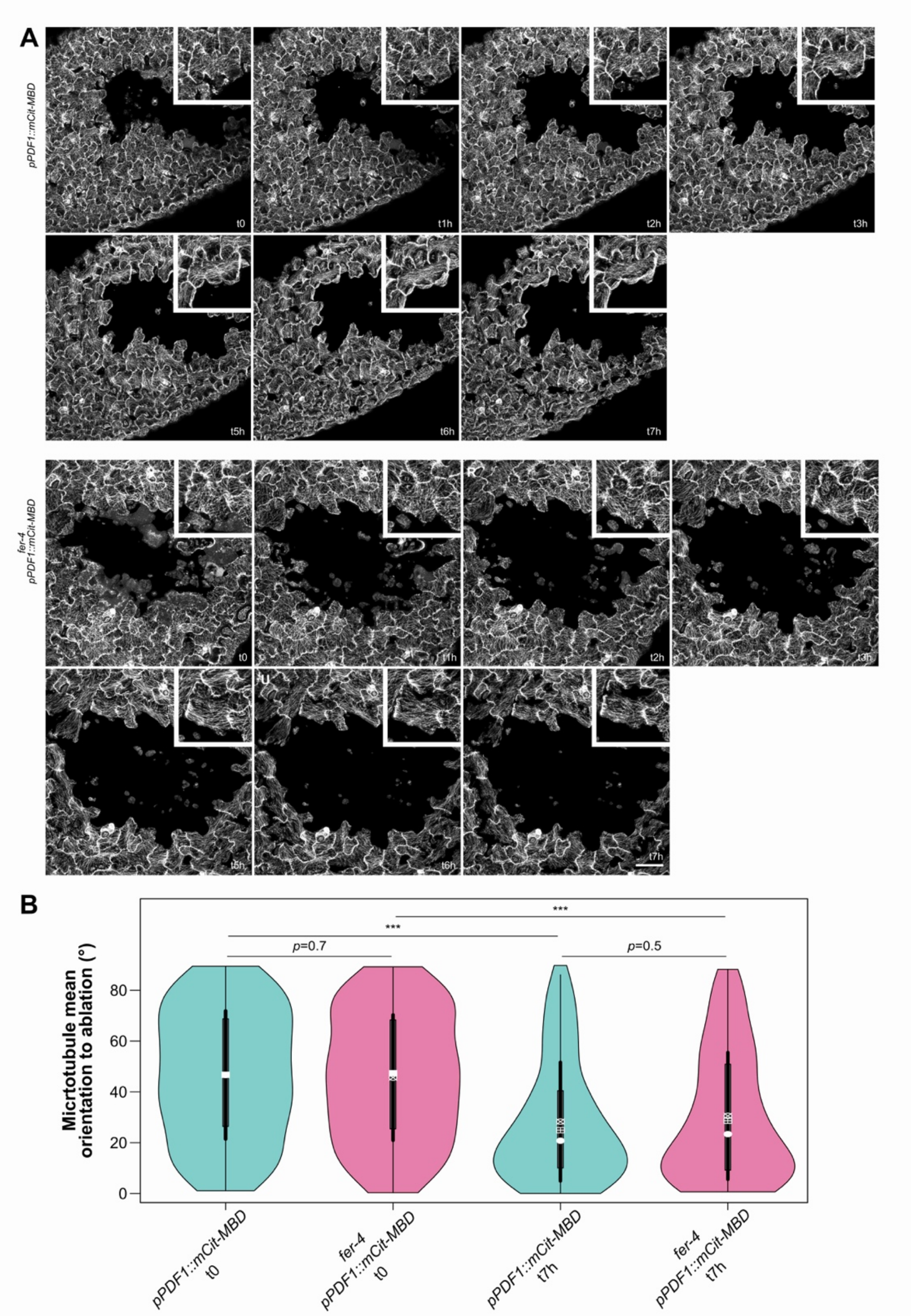


**Supplementary Figure 4.** Ablations (full kinetics) on 2.5% agar medium

**A.** Representative confocal images of *pPDF1::mCit-MBD* (top) and *fer-4 pPDF1::mCit-MBD* (bottom) pavement cells, from seedlings grown on a medium containing 2.5% agar, immediately after an ablation (t0) and 1 to 7 hours later. Scale=50µm. **B.** Orientation to the ablation (violin plot) of cortical microtubule arrays and *p*-values (*p*) of Wilcoxon-Mann-Whitney tests in cells surrounding the ablation site in *pPDF1::mCit-MBD* and *fer-4 pPDF1::mCit-MBD* pavement cells, immediately after an ablation (t0) and 7 hours later (t7h).

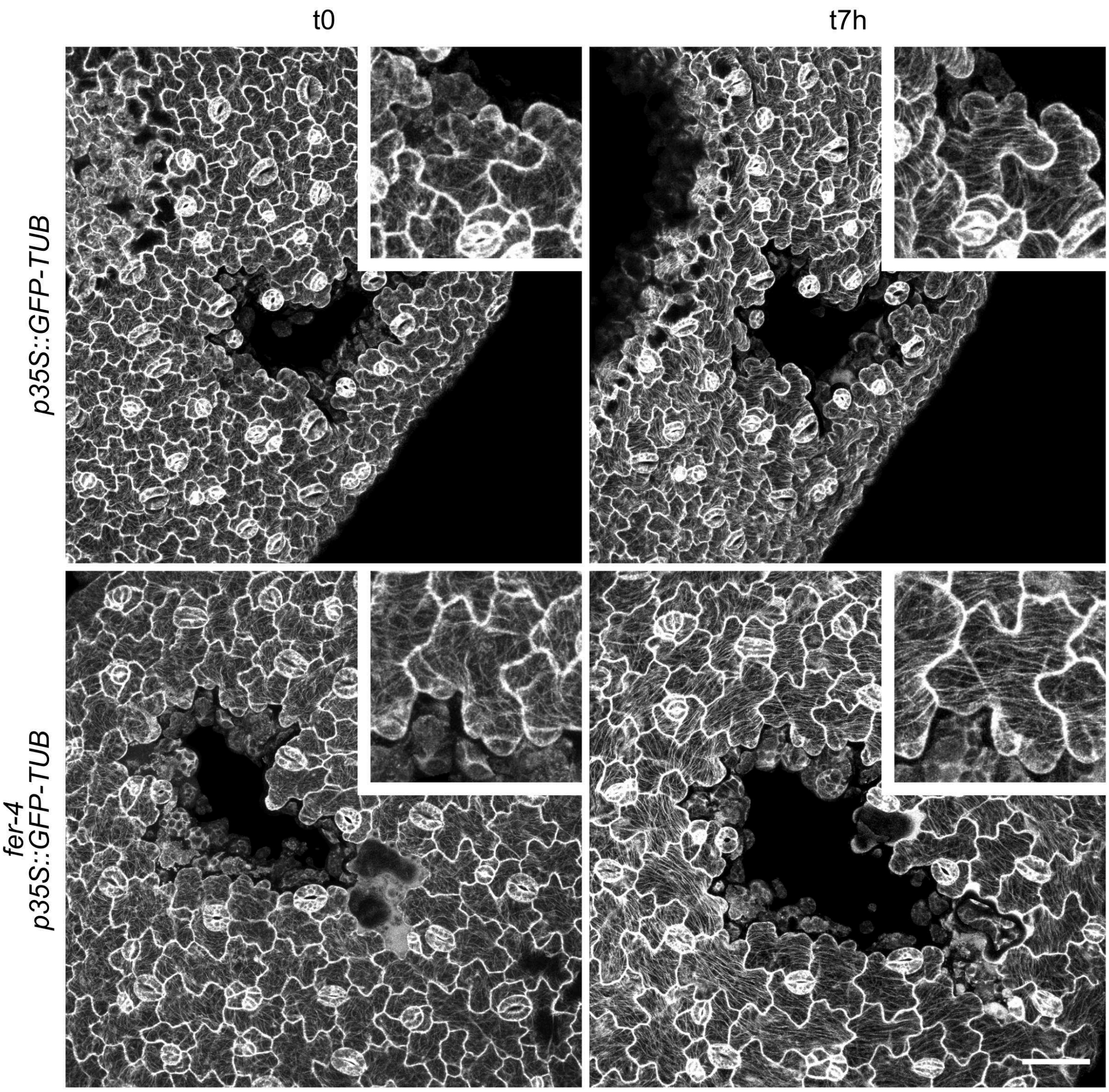


**Supplementary Figure 5.** Microtubule response after ablation using the GFP-tubulin reporter

Representative confocal images of *p35S::GFP-TUB* (top) and *fer-4 p35S::GFP-TUB* (bottom) pavement cells from seedlings grown on a medium containing 2.5% agar, immediately after an ablation (t0) and seven hours later (t7h). Scale=50µm.

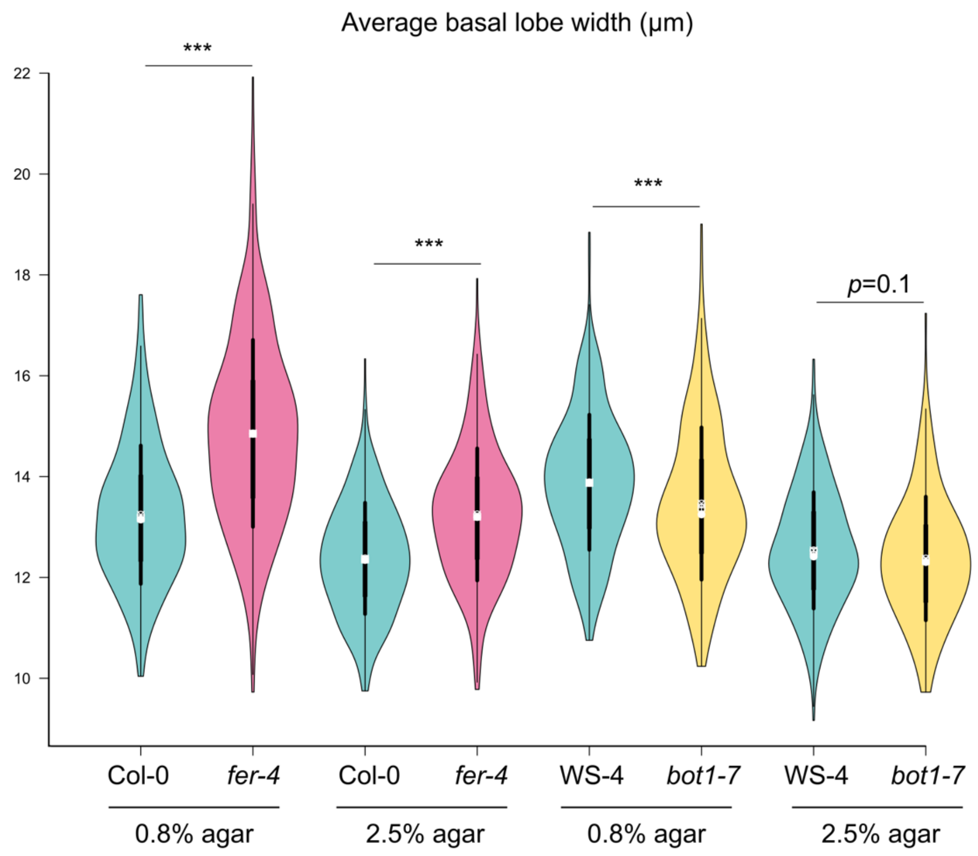


**Supplementary Figure 6.** Basal lobe width according to genotype and growth conditions.

Basal lobe width (violin plot) of pavement cells and *p*-values (*p*) of Dunn tests for the WT (*Col-0, WS-4*), katanin mutant (*bot1-7*), and *fer-4*. Seedlings were grown on 0.8% or 2.5% agar. Data from Figure 5A (0.8% agar) are reproduced here to allow a full comparison of the results.
